# Opposing Microtubule Motors Control Motility, Morphology, and Cargo Segregation During ER-to-Golgi Transport

**DOI:** 10.1101/001743

**Authors:** Anna K. Brown, Sylvie D. Hunt, David J. Stephens

**Affiliations:** Cell Biology Laboratories, School of Biochemistry, Medical Sciences Building, University of Bristol, BS8 1TD

**Author notes:** NÉE ANNA K. TOWNLEY.

**Keywords:** Microtubule Motor, Dynein, Kinesin, Endoplasmic Reticulum, Golgi, Secretory Cargo Trafficking

## Abstract

We recently demonstrated that dynein and kinesin motors drive multiple aspects of endosomal function in mammalian cells. These functions include driving motility, maintaining morphology (notably through providing longitudinal tension to support vesicle fission), and driving cargo sorting. Microtubule motors drive bidirectional motility during traffic between the endoplasmic reticulum (ER) and Golgi. Here, we have examined the role of microtubule motors in transport carrier motility, morphology, and domain organization during ER-to-Golgi transport. We show that consistent with our findings for endosomal dynamics, microtubule motor function during ER-to-Golgi transport of secretory is required for motility, morphology of, and cargo sorting within vesicular tubular carriers *en route* to the Golgi. Our data are consistent with previous findings that defined roles for dynein-1 and kinesin-1 (KIF5B) and kinesin-2 in this trafficking step. Our high resolution tracking data identify some intriguing aspects. Depletion of kinesin-1 reduces the number of motile structures seen which is in line with other findings relating to the role of kinesin-1 in ER export. However, those transport carriers that were produced had a much greater run length suggesting that this motor can act as a brake on anterograde motility. Kinesin-2 depletion did not significantly reduce the number of motile transport carriers but did cause a similar increase in run length. These data suggest that kinesins act as negative regulators of ER-to-Golgi transport. Depletion of dynein not only reduced the number of motile carriers formed but also caused tubulation of carriers similar to that seen for SNX-coated early endosomes. Our data indicated that the previously observed anterograde-retrograde polarity of transport carriers in transit to the Golgi from the ER is maintained by microtubule motor function.

## Introduction

Endoplasmic reticulum (ER)-to-Golgi transport is required for the delivery of newly synthesized secretory cargo to the Golgi apparatus. Following the concentration of cargo and budding of carriers at the ER, cargo accumulates in the ER-Golgi intermediate compartment (ERGIC) from which it is then transported to the Golgi. Transport of cargo in small vesicular tubular transport carriers between the ER and Golgi along microtubules ensures vectorial delivery of cargo in an efficient manner (Ho et al., 1989; Lippincott-Schwartz et al., 1990; Thyberg and Moskalewski, 1985). The link to microtubules is established at the point of ER export through direct interaction of the Sec23 subunit of the COPII coat complex with the p150^Glued^ subunit of dynactin (Watson et al., 2005). It is not known how the ERGIC is itself linked to microtubules but considerable data have shown that cytoplasmic dynein-1 is required for the movement of cargo in VTCs to the Golgi (Burkhardt et al., 1997; Palmer et al., 2009; Presley et al., 1997). Other work has identified roles for kinesin-1 in ER export and in retrograde transport from the Golgi back to the ER (Lippincott-Schwartz et al., 1995).

Kinesin-2 also acts in retrograde traffic from the Golgi (Stauber et al., 2006). Recently, we used RNAi-mediated depletion of individual microtubule motor subunits to define their function in the endosomal system (Hunt et al., 2013). Using GFP-tagged sorting nexins (SNXs) and total internal reflection fluorescence (TIRF) microscopy to maximize signal-to-noise ratios, we found that specific combinations of microtubule motors were coupled to individual sorting SNX-coated membrane domains.

We showed that these specific combinations of motors direct not only the motility of these endosomal domains but also dictate their morphology and the segregation of cargo within them. Our data are consistent with models in which longitudinal tension, generated by the motor acting on a membrane (Roux et al., 2002), aids in the scission event that generates small transport carrier from larger compartments (reviewed in (Stephens, 2012)). This longitudinal tension could form part of a geometric sorting process driving cargo segregation into distinct functional domains (Mayor et al., 1993). Since ER-to-Golgi transport, like early endosomal motility, is known to use cytoplasmic dynein-1 in combination with kinesin-1 and kinesin-2, we chose to apply the same technique to the transport of secretory cargo from the ER to the Golgi. We chose tsO45-G-GFP for this process as this viral glycoprotein is widely used model cargo and provides a means to initiate cargo trafficking through temperature-controlled. At 39.5°C tsO45-G-GFP is retained in the ER; at 32°C it folds and rapidly exits the ER using the same machinery as other endogenous cargo. While a non-homologous system, the benefits of being able to image transport in a controlled way outweigh the limitations. Furthermore, the increase in signal to noise ratio using of TIRF microscopy allowed us to use this probe at a relatively low expression. We use TIRF microscopy here not to provide a selective illumination of events very close to the plasma membrane but instead with an increased penetration depth to facilitate this increase in signal-to-noise.

Previous work has shown that tsO45-G-GFP is trafficked in vesicular tubular clusters to which the COPI coat complex is recruited. COPI directs retrograde sorting of membrane proteins from the Golgi to the ER and a widely accepted model is that this retrograde recycling is initiated at the ERGIC and continues during transit of cargo to the Golgi. An interesting feature of this process is that COPI localizes to the retrograde domain of these carriers while secretory cargo such as tsO45-G-GFP accumulates in the Golgi-facing or anterograde domain (Shima et al., 1999). This domain-based segregation of cargo is reminiscent of that seen in the endosomal pathway.

Here we have taken this same approach of motor subunit depletion using RNAi coupled with TIRF imaging to examine the role of microtubule motors in the dynamics of transport carriers moving from the ER to the Golgi.

## Results

We used TIRF imaging of cells expressing the temperature sensitive form of the vesicular stomatitis virus glycoprotein (ts045-G) tagged with GFP. This enables controlled release of this model secretory cargo from the ER such that we can unequivocally identify structures during the early stages of transit to the Golgi (<12 minutes after release of the temperature block). From analysis of many experiments, we determined that the number of detectable ts045-G-GFP-positive structures was decreased following either dynein-1 or kinesin-1 depletion (but not that of other motors) the proportion of motile structures was not changed. This decrease in formation of VTCs is consistent with our previous work (Gupta et al., 2008). Motor protein depletion was achieved using siRNA transfection and validated by immunoblotting periodically through the project (see also (Hunt et al., 2013; Palmer et al., 2009)).

Figure 1 shows maximum intensity projections of the transport of tsO45-G-GFP starting 12 minutes after the shift to 32°C. These are derived from the movie sequences shown in the associated movie files. Two versions of each movie are included, one playing back at real time, and another playing back at 3x speed. In control cells (Fig. 1A–C, Movie 1A and Movie 1B) many tracks are evident that translocate towards the Golgi. We then performed the same assay in cells that had previously been depleted of individual motor subunits prior to expression of tsO45-G-GFP. Suppression of dynein heavy chain 1 (DHC1) resulted in fewer visible carriers translocating to the Golgi (Fig. 1D, Movie 2A and Movie 2B). An immediate feature of the maximum intensity projections is that the length of individual tracks is much shorter (Fig. 1E, F and see below). Many small punctate carriers were still visible which we attribute to the incomplete loss of DHC1 expression in these experiments. Our interpretation is therefore based on changing the balance of cellular motor subunits and not the presence or absence of any individual component. In our own work, we have previously defined a role of the light intermediate chain 1 (LIC1) of cytoplasmic dynein 1 in Golgi organization and ER-to-Golgi transport (Palmer et al., 2009). Here we found that depletion of LIC1 led to a similar but less dramatic phenotype to depletion of DHC1 (Fig. 1G, Movie 5A and Movie 5B). Notably an increase in the number of static structures and those only moving a very short distance was evident (Fig. 1H, enlarged in I). KIF5B had previously been implicated in the export of cargo from the ER (Gupta et al., 2008) and kinesin-2 has also been implicated in Golgi-to-ER retrograde trafficking (Stauber et al., 2006). In many other systems motors of opposing polarity are essential for movement in either direction (Ally et al., 2009). Here we found that in contrast to depletion of dynein subunits where motility was reduced, kinesin-1 depletion led to increased movement of tsO45-G-GFP along continuous tracks that were substantially longer than seen in controls (Figure 1 J–L, Movie 3A and Movie 3B). A similar result, although less noticeable, was seen following depletion of kinesin-2 (achieved by targeting the KAP3 accessory subunit of this motor, Figure 1M–O, Movie 4A and Movie 4B).

**Figure 1.**
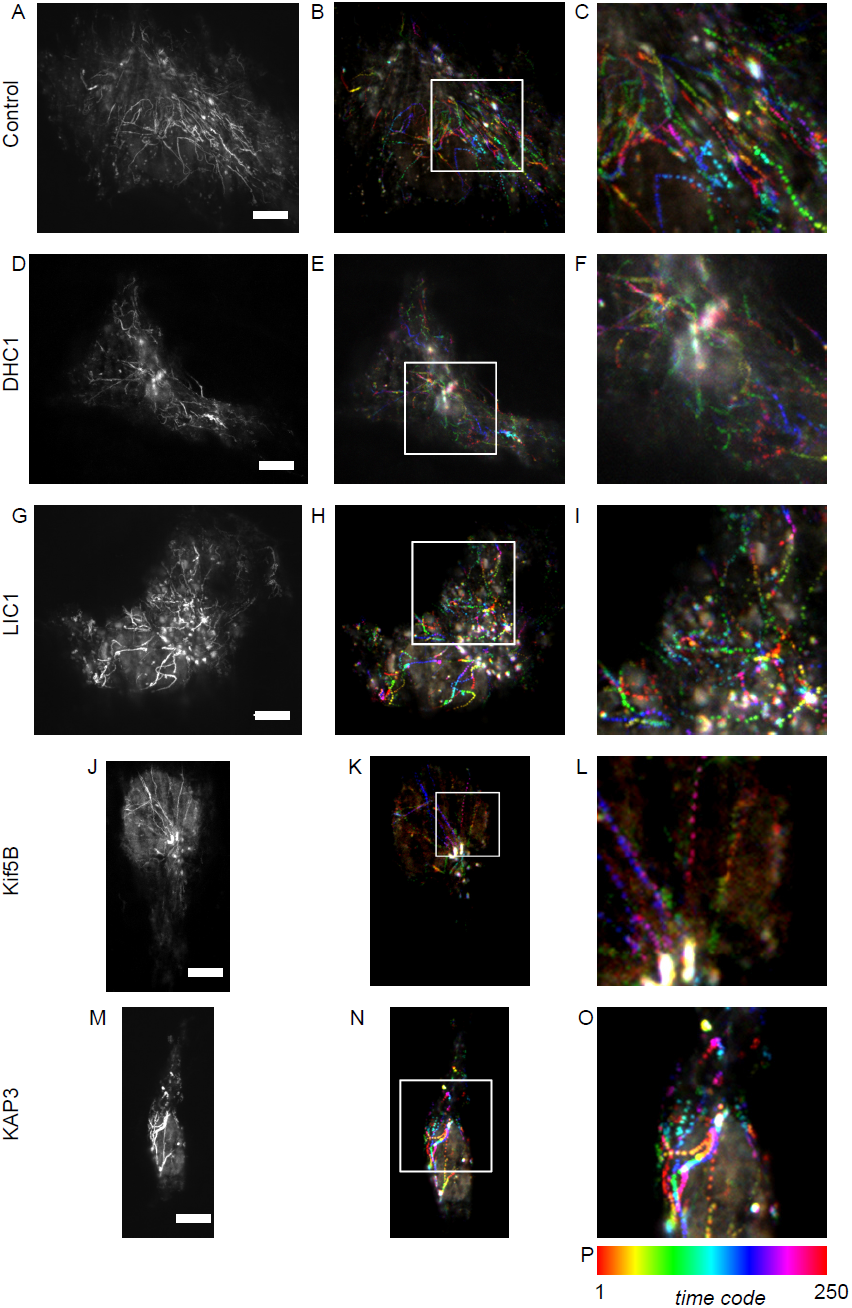
Projections of time lapse sequence from TIRF imaging of tsO45-G-GFP translocation. Image sequences are comprised of 1000 frames acquired at 12.04 frames per second. (A, D, G, J, M) Maximum intensity projections generated from all 1000 frames. (B, E, H, K, N) enlarged in (C, F, I, L, O) show colour coded time projections of every fourth frame from these sequences (332 ms intervals, 250 frames total). (P) Shows the colour coding for time projections. Scale bars (all frames) = 10 μm.

Quantification of these data is presented in Figure 2. Depletion of either DHC1 or KIF5B significantly reduced the number of motile transport carriers (defined as those moving >5 μm) that we observed (Fig. 2A). This likely reflects the known function of these motors in ER export (Gupta et al., 2008) and transport carrier motility (Presley et al., 1997). This contrasts with our findings in relation to endosomal motility where opposing motors are both required to drive motility (Hunt et al., 2013) and peroxisome motility (Ally et al., 2009). We then measured the movement of objects based on track length from maximum intensity projections from time sequences of 50 seconds duration. In all cases, we found that the speed of individual moving structures was the same regardless of gene depletion using RNAi, ranging from 0.2-3.1 μm.sec^−1^. Fig. 2B shows the spread of track lengths for individual structures measured from maximum intensity projections of time sequences. In control cells a range of track lengths between 0 and 160 μm was measured. Depletion of DHC1 led to a significant reduction in track length; the mean track length was also decreased on suppression of LIC1 but this difference was not statistically significant. Of note, the longest tracks in this case are those that are lost. We conclude from these plots, and the image data in Fig. 1, that LIC1 suppression was less effective at inhibiting motility of these structures perhaps due to partial redundancy with LIC2 or due to incomplete suppression. In this study, we have not analyzed cells following LIC2 suppression only as our previous work strongly implicated LIC1 in Golgi-related functions of dynein (Palmer et al., 2009). Somewhat surprisingly, depletion of either kinesin-1 or kinesin-2 resulted in a statistically detectable increase in track length (Fig. 2, green asterisks indicate a statistically detectable increase in track length). The statistical tests are consistent with the visual impression of the data shown in Fig. 1 that kinesin-2 depletion is less effective than that of KIF5B. These data imply that kinesins act against dynein in this system to attenuate motility of ER-to-Golgi transport carriers.

**Figure 2.**
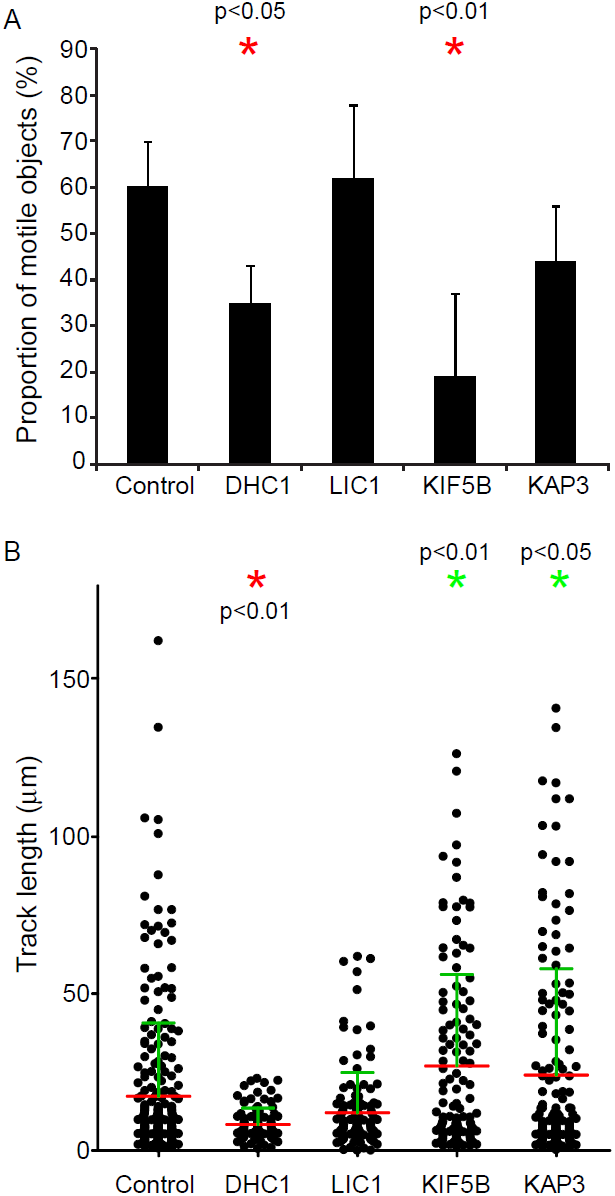
(A) Proportion of moving objects as a percentage of the total puncta detected. Statistical significance was tested using an ANOVA with a Dunnett’s post-hoc test. (B) Quantification of time-lapse imaging. Images were processed to generate maximum intensity projections as shown in Figure 1. Track lengths were then automatically measured in Volocity and plotted. Each spot represents an individual particle. Measurements are derived from 3 independent experiments, 3-8 cells in each experiment, total of 75 objects. Statistical significance was tested using an ANOVA with a Dunnett’s post-hoc test. Red asterisks indicate statistically detectable decrease in track length; green asterisks represent a statistically detectable increase in track length.

From our work on the endosomal system, we determined that microtubule motors were required to maintain the morphology of endosomes (Hunt et al., 2013). Depletion of individual motor subunits led to tubulation of SNX-labelled endosomal compartments. We have previously documented a similar effect on early ER-to-Golgi transport carriers following KIF5B depletion (Gupta et al., 2008). Here, in our TIRF time-lapse sequences tracking the movement of tsO45-G-GFP from the ER to the Golgi we observed a similar, although less dramatic, increase in tubulation following DHC1 suppression (Fig. 3, multiple examples are shown that are taken from the time sequence in Fig 1D and movie 2). Tubules were occasionally seen in control movies (1-2 per cell in the 50 second time sequences acquired) but were more obvious in DHC1-depleted cells (3-10 per cell). Tubules were typically short (< 5 μm), highly motile, and were occasionally branched (Fig. 3A). Other work has shown that tubulation of ER-to-Golgi carriers is linked to the level of cargo expression (Simpson et al., 2006). Analysis of fluorescence intensity in the data sets described here indicates that the increased tubulation seen here is due to DHC1 suppression and not an increase in tsO45-G-GFP expression in these selected cases (not shown). These data are consistent with work showing that tubular transport intermediates are dependent on an intact microtubule network (Simpson et al., 2006), and that describing the formation of highly dynamic ERGIC tubules following motor depletion (Tomas et al., 2010).

**Figure 3.**
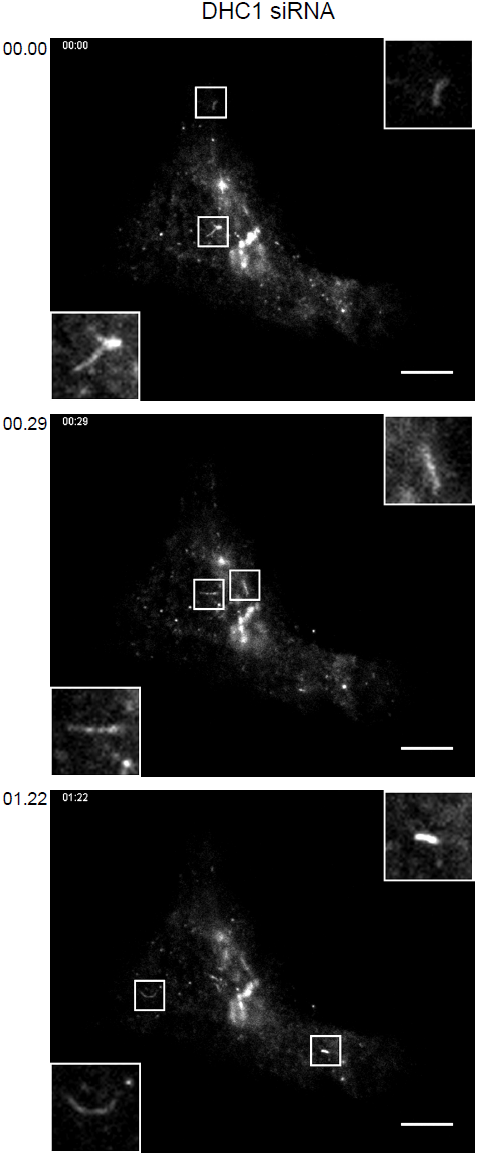
Increased tubulation of ER-to-Golgi carriers following suppression of DHC1 expression. Examples are taken from the time lapse movie sequence associated with Figure 1D. Boxed regions show 3x enlargements. Time from the start of the image sequence is shown (mins:secs) Bars = 10 μm.

Analysis of the role of motor subunits in the endosomal system also led us to define a role in segregation of endocytic cargo into distinct functional domains (Hunt et al., 2013). Previous work has shown that ts045-G-GFP localizes to an anterograde (i.e. Golgi facing) domain of the transport intermediate during its transit to the Golgi (Shima et al., 1999). The COPI complex provides a marker of the retrograde domain. We used these markers in fixed cells, 12 minutes after temperature shift to 32°C to ask whether dynein and kinesin motors are required to maintain this domain organization.

Objects were measured by visual inspection with the Golgi defined as the centre of the brightest cluster of objects (denoted with the letter “G” in Fig. 4A). Here it is important to note that depletion of DHC1 causes significant disruption to Golgi structure, resulting in the presence of multiple small Golgi elements throughout the cytoplasm. This complicates analysis here likely leading to an under estimation of the degree of COPI-association with bona fide ER-to-Golgi transport carriers and likely an inaccurate assessment of their polarity. Regardless, we include these data for completeness. Fig. 4A shows and example of these image data from control cells (GL2 siRNA transfected). Green arrows indicate the position of tsO45-G-GFP and red arrows the position of COPI. Insets 1, 2, and 4, show a polarization of the structure with the green domain closer to the Golgi (indicated by the letter “G”). Inset 3 shows an object of the opposite polarization with the COPI-positive domain facing the Golgi. We measured two parameters from these data: the number of tsO45-G-GFP puncta that were positive for COPI, and the polarization of these structures with regard to the Golgi. Fig. 4B shows the proportion of tsO45-G-GFP structures that were positive for COPI. Gray bars indicate the mean. Depletion of LIC1, KIF5B, or KAP3 did not affect the association of COPI with these structures. This is consistent with the known mechanisms of recruitment of COPI to membranes and is suggests that these motor subunits are not required for the recruitment of COPI. While a statistically significant decrease in COPI association is seen following DHC1 depletion, we attribute this to the fragmentation of the Golgi apparatus in these cells and the fact that even at 12 minutes after temperature shift, there is a substantial proportion of tsO45-G-GFP in the early Golgi. In contrast, when we measured the orientation of tsO45-G-GFP and COPI positive structures, we found that depletion of any of the four motor subunits resulted in a decrease in the proportion of correctly polarised objects (Fig. 4C). In particular, depletion of LIC1 or of KAP3 resulted in a statistically detectable decrease in this measure (Fig 4C, red asterisks). The microtubule network was not perturbed significantly in these cells (see (Gupta et al., 2008; Hunt et al., 2013; Palmer et al., 2009)) and thus, we conclude that these motors are required to either generate or maintain this domain organization.

We then sought to define whether there was any obvious defect in ER-to-Golgi cargo trafficking. For this we chose to analyse the distribution of KDEL-bearing proteins which under normal circumstances are retained in the ER by active retention and retrieval mechanisms. We reasoned that a possible defect in COPI-dependent retrieval following motor subunit depletion might manifest itself as an increase in anterograde transport of KDEL-bearing cargoes. Immunoblotting (Fig. 4D) and immunofluorescence (Fig. 4D, showing images of control and KIF5B siRNA transfected cells) of KDEL-bearing proteins revealed no differences in steady-state localization in control versus motor subunit depleted cells.

## Discussion

The importance of the microtubule motor system to mammalian cell organization and function is unquestionable. Our knowledge of the mechanochemistry of these enzymes, as well as the ways in which they are targeted to specific intracellular cargoes is increasing rapidly. In this study, we have extended our work examining the role of individual motor subunits in the endocytic pathway to the biosynthetic secretory pathway. This has highlighted several common features and some notable differences. In general terms we can conclude that in both the endocytic and biosynthetic systems, microtubule motors act not only in the motility of these small vesicular and tubular carriers but also in maintaining their structure, and in maintaining (and possibly generating) the functionally distinct domains within these complex structures.

Our data also highlight cross-talk between motors of opposing polarity, dynein and the kinesins, in ER-to-Golgi transport. Suppression of KIF5B expression results in longer linear tracks of tsO45-G-GFP suggesting that it normally acts as a negative regulator of run length, probably through opposing the minus-end directed force of dynein. The normal balance of opposing motors that frequently engage in a tug-of-war (Derr et al., 2012) is likely imbalanced by our KIF5B gene depletion experiments such that we bias movement towards the minus end. That we observe this effect more robustly with KIF5B depletion than KAP3 depletion suggests that dynein-1 opposing kinesin-1 is the most common tug-of-war in this ER-to-Golgi trafficking step. While we can clearly identify these changes in transport carrier motility following motor subunit depletion we have been unable to define any functional consequence of this. KIF5B depletion does slow the overall rate of cargo trafficking (Gupta et al., 2008) but only at later stages release of tsO45-G-GFP from the ER. The Golgi remains *cis trans* polarized in the same way that it is in control cells, we do not observe any gross defect in retrograde trafficking of KDEL tagged cargo in KIF5B depleted cells (which used uses the COPI system in concert with the KDEL receptor (Lewis and Pelham, 1990; Munro and Pelham, 1987)).

It is possible, even probable that defects in these motor-dependent trafficking events on manifest themselves as gross phenotypic consequences in cell types that have a significant dependence on long range microtubule-based transport. An obvious example would be motor neurons and this is of course consistent with the fact that many mutations in dynein subunits, kinesins, and their accessory molecules result in severe neurological defects (recently reviewed in (Franker and Hoogenraad, 2013)). We have not undertaken any experiments to address this at present.

The identity of the molecular linkers between these motors and the membranes that they carry are still not well defined. Recent work has identified the Golgi matrix protein golgin-160 as a key determinant of dynein localization to the Golgi (Yadav et al., 2012). Whether golgin-160 is the specific, or indeed the only, dynein anchor in ER-to-Golgi transport of anterograde cargo remains unclear. This is an important question as the molecular basis for trafficking through the early secretory pathway is still poorly understood. It is know that COPII mediates the export of cargo from the ER. COPII-derived vesicles then either fuse together homotypically or fuse with a pre-existing ER-Golgi intermediate compartment (ERGIC). The subsequent transport step remains something of a mystery. Is there a budding event from the ERGIC or are large parts of this organelle separated from the underlying structure prior to transport to the Golgi? This latter option could clearly involve the application of force by motor proteins, notably dynein, in an analogous way to the role of kinesin at the *trans*-Golgi network (Polishchuk et al., 2003). Analysis of these options is complicated by the fact that perturbation of the ERGIC often results in an inhibition of COPII-derived budding. An example is the use of brefeldin A and other GBF1 inhibitors that while acting on recruitment of COPI through inhibiting Arf activation result in accumulation of secretory cargo, not in the ERGIC, but in the ER (Lippincott-Schwartz et al., 1989). Clearly the identification of factors that control motor recruitment and activation on ERGIC and transport carrier membranes will help solve these questions.

## Materials and Methods

All reagents were purchased from Sigma-Aldrich (Poole, UK) unless otherwise stated.

### Growth of Culture Cells

Human telomerase immortalized retinal pigment epithelial cells (hTERT-RPE1) were maintained in DMEM-F12 supplemented with 10% FCS (Life Technologies, Paisley, UK) containing supplemented 1% L-glutamine and 1% essential amino-acids. At 24 hours prior to the start of the experiments, cells were seeded onto 35 mm glass bottom dishes (MatTek, Ashland, MA).

### Source of Antibodies and Other Reagents

Monoclonal rat anti-human tubulin (ab6160) and polyclonal rabbit anti-KIF5B (ab5629) were from Abcam (Cambridge); polyclonal rabbit anti-human kinesin-2 (kindly provided by Isabelle Vernos, Barcelona, Spain); polyclonal rabbit anti-human lamin A/C (#2032) was from Cell Signaling Technology (Danvers, MA). Secondary antibodies were from Jackson ImmunoResearch Laboratories (West Grove, PA). Anti-KDEL antibody (clone 10C2) was from Abcam.

### Small Interfering RNA Transfection

Cells were siRNA-transfected by calcium phosphate method at 3% CO_2_ (Chen and Okayama, 1988). The medium was changed 20 hours after transfection and cells were washed with PBS and were incubated for 72 hours at 37°C at 5% CO_2_ with fresh supplemented media. SiRNA duplexes were designed using online algorithms of, and subsequently synthesized by, MWG-Eurofins. BLAST search was performed for these duplexes against the non-redundant database to determine their specificity. Lamin A/C or luciferase GL2 were depleted as targeted controls. Sequences used were as follow: DHC1-a ACA UCA ACA UAG ACA UUC ATT; DHC1-b CCA AGC AGA UAA GGC AAU ATT; KIF5B-a UGA AUU GCU UAG UGA UGA ATT; KIF5B-b UCA AGU CAU UGA CUG AAU ATT; KAP3-a CUU GAC CAU UCC AGA CUU ATT; KAP3.b GCU CUG UGU AUG AAU AUU ATT; DLIC1-a AGA UGA CAG UGU AGU UGU ATT; DLIC1-b GAA CAU GAC UAC AGA GAU GTT; Lamin A/C CUG GAC UUC CAG AAG AAC ATT; GL2 CGU ACG CGG AAU ACU UCG AUU TT.

### Transient Plasmid Expression

Transient expression of tsO45-G-GFP was done using Lipofectamine^™^ 2000 as per manufacturer’s instructions.

### Immunolabelling

Medium was removed and cells were subsequently washed with PBS. Cells were then fixed using cold methanol for 4 minutes at -20°C. After two washes in PBS, cells were blocked using a 3% bovine serum albumin (BSA) in PBS for 30 minutes at room temperature. Cells were incubated with the primary antibodies (anti-beta’-COP rabbit polyclonal “BSTR”) in 3% BSA for 1 hour at room temperature. Three washes with PBS of 5 minutes each at room temperature were done before incubating the secondary antibody. The latter was diluted at 1:400 in PBS and incubated 1 hour at room temperature. Finally, after three washes in PBS (5 minutes each at room temperature) and an optional nuclear staining with DAPI (4,6-Diamidino-2-phenylindole, Molecular Probes, diluted at 1:5000 in distilled water) for 3 minutes at room temperature, cells were gently rinsed twice in PBS.

### Immunoblotting

For immunoblots to validate siRNA efficacy (not shown but for examples see (Gupta et al., 2008; Hunt et al., 2013; Palmer et al., 2009)), cells were lysed and samples were separated by SDS-PAGE followed by transfer to nitrocellulose membranes; primary antibodies were detected using HRP-conjugated secondary antibodies (Jackson ImmunoResearch, West Grove, PA) and enhanced chemiluminescence (ECL, GE Healthcare, Cardiff, United Kingdom). For analysis of KDEL protein secretion (Fig. 4) blots were developed using fluorophore conjugated secondary antibodies and an Odyssey Sa imaging system (LI-COR Biosciences, Cambridge, UK).

**Figure 4.**
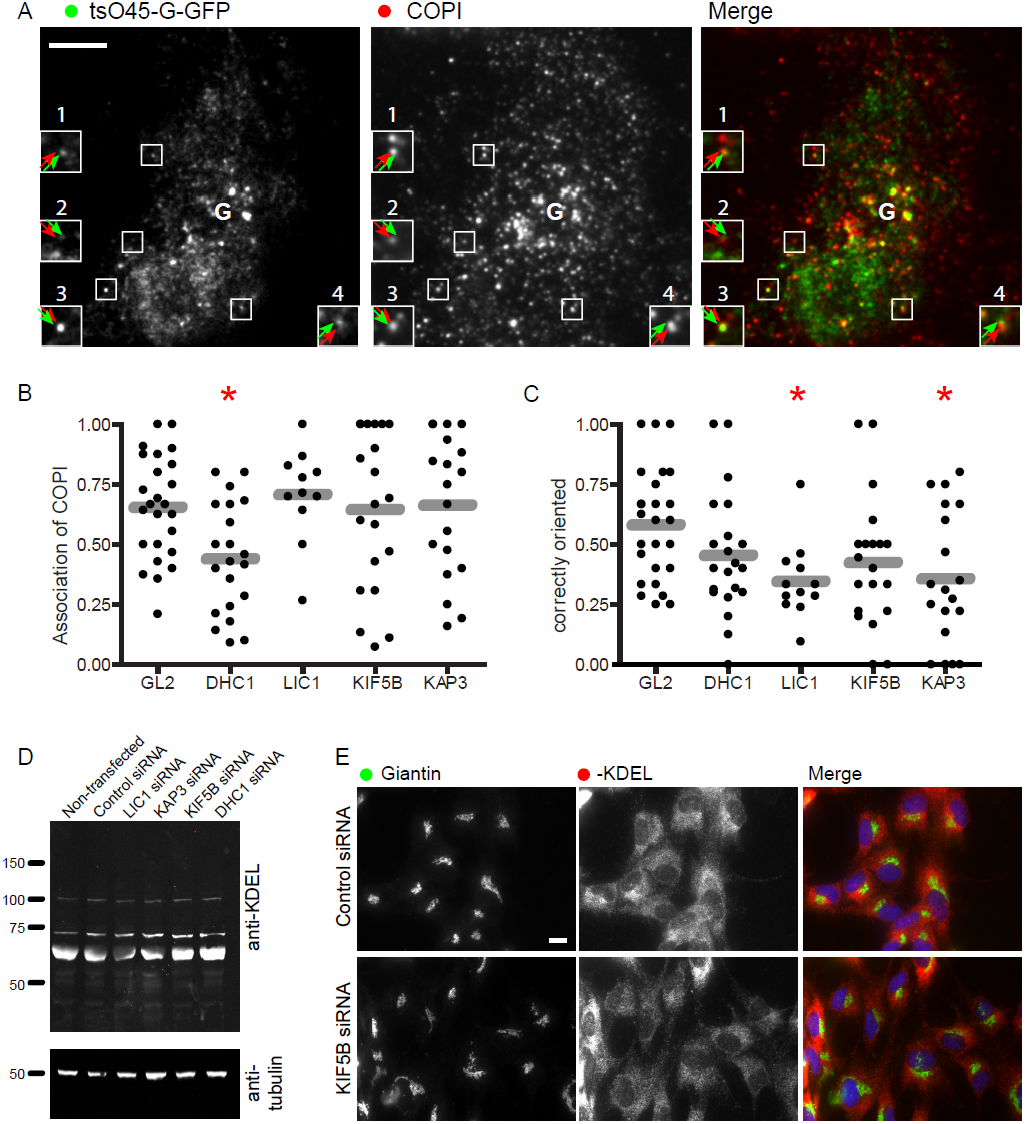
Motor subunit depleted cells expressing tsO45-G-GFP were fixed and immunolabelled to detect COPI (β’-COP). (A) An example of a control (GL2 siRNA transfected) cell. Bar = 10 μm. (B, C) Image quantification was performed manually to determine (B) the number of tsO45-G-GFP puncta with associated COPI labelling and (C) the polarity of these structures with respect to the Golgi apparatus. “Correctly oriented” structures are those with the COPI-positive domain facing away from the Golgi and the tsO45-G-GFP-positive domain facing towards the Golgi. G indicates the user-defined point of the Golgi apparatus. Asterisks in B and C in dictate those cases for which a statistically detectable difference was found using ANOVA with Dunnett’s post-hoc test. (D) RPE1 cells were transfected with siRNA against the various motor proteins for 72 hours. The cells were then lysed and after a protein assay equal amounts of protein analyzed by immunoblotting with an anti-KDEL or an anti-α-tubulin antibody. (E) Immunofluorescence labelling of control or KIF5B depleted cells with antibodies directed against giantin or –KDEL bearing proteins. Bar (all panels) = 10 μm.

### TIRF Imaging

Cells were kept at 37°C and at 5% CO2. Media was discarded and replaced with pre-warmed imaging medium (Life Technologies) just prior imaging. The TIRF system used was a Leica AM-TIRF MC (multi colour) associated with a Leica DMI 6000 inverted microscope. Images were acquired with an oil immersion TIRF objective with 100x 1.46 numerical aperture and were acquired at the rate 12.04 frames per second (i.e. the interval between images was 83 ms) using a Hamamatsu C9100-13 EMCCD camera. Each image was 512 by 512 pixels with no additional binning. 488 nm and 561 nm lasers were used for excitation. Data was processed using Volocity software (Perkin Elmer) and Adobe Illustrator CS (Adobe). TIRF Imaging was used in all assays, including of fixed cells in Fig. 4. The penetration depth used was set at 150 nm to optimise the ratio signal / noise. For motor-dependent endosomal sorting assays, thousand frames were acquired. Colocalization studies were also performed on live cells to keep the tubular structures intact. Image brightness and contrast was adjusted in ImageJ/Fiji and the same processing was applied to all image sets within each experiment.

### Processing and Quantification of Image Data and Statistical Analysis

Maximum intensity projections in Figure 1A, D, G, J, and M were done using Fiji. Colour coding of time sequences was achieved using an ImageJ plug-in developed by Kota Miura (EMBL Heidelberg) which is available at: http://cmci.embl.de/downloads/timeseriescolorcoder and within the Fiji implementation of ImageJ (Schindelin et al., 2012). For these data, the first image of each sequence was extracted from the sequence and then subtracted from the remaining data set to reduce background prior to running the “time series color coder” plugin.

Track lengths were measured automatically in Volocity (version 5.4.2, Perkin Elmer) after thresholding to detect only lines >1 μm. Data in Fig. 2 come from measuring >750 objects from 3 independent experiments.

Representative images are shown, all experiments were repeated independently at least three times each. Samples were compared using ANOVA with Dunnett’s post-hoc test. All images were prepared with Adobe Photoshop CS and Adobe Illustrator CS3.

## Acknowledgements

We would like to thank the Wolfson Foundation, MRC, and University of Bristol for establishing and maintaining the University of Bristol Bioimaging Facility and Mark Jepson, Alan Leard, and Katy Jepson for their help with training.

### Contributions

A.K.T. performed experimental work, analyzed the data and helped draft the manuscript; S.D.H. provided significant support to assay development and data analysis; D.J.S. conceived and directed the project, analyzed data, and wrote the manuscript.

### Funding

This work was funded by the UK Medical Research Council (grant number G0801848). The authors declare no competing financial interests.

## Supplementary Material: Movies

All movies show TIRF imaging of tsO45-G-GFP 12 minutes after temperature shift of cells to 32°C. Cells were previously transfected with siRNA duplexes as indicated.

Movie 1A: GL2-transfected cells. Playback at real time. See Figure 1A.

Movie 1B: GL2-transfected cells. Playback at 3x real time. See Figure 1A.

Movie 2A: DHC1-transfected cells. Playback at real time. See Figure 1D.

Movie 2B: DHC1-transfected cells. Playback at 3x real time. See Figure 1D.

Movie 3A: KIF5B-transfected cells. Playback at real time. See Figure 1G.

Movie 3B: KIF5B-transfected cells. Playback at 3x real time. See Figure 1G.

Movie 4A: KAP3-transfected cells. Playback at real time. See Figure 1J.

Movie 4B: KAP3-transfected cells. Playback at 3x real time. See Figure 1J.

Movie 5A: LIC1-transfected cells. Playback at real time. See Figure 1M.

Movie 5B: LIC1-transfected cells. Playback at 3x real time. See Figure 1M.

## References

Ally, S., A.G. Larson, K. Barlan, S.E. Rice, and V.I. Gelfand. 2009. Opposite-polarity motors activate one another to trigger cargo transport in live cells. The Journal of cell biology. 187:1071–1082.

Burkhardt, J.K., C.J. Echeverri, T. Nilsson, and R.B. Vallee. 1997. Overexpression of the dynamitin (p50) subunit of the dynactin complex disrupts dynein-dependent maintenance of membrane organelle distribution. The Journal of cell biology. 139:469–484.

Chen, C.A., and H. Okayama. 1988. Calcium phosphate-mediated gene transfer: a highly efficient transfection system for stably transforming cells with plasmid DNA. BioTechniques. 6:632–638.

Derr, N.D., B.S. Goodman, R. Jungmann, A.E. Leschziner, W.M. Shih, and S.L. Reck-Peterson. 2012. Tug-of-war in motor protein ensembles revealed with a programmable DNA origami scaffold. Science. 338:662–665.

Franker, M.A., and C.C. Hoogenraad. 2013. Microtubule-based transport - basic mechanisms, traffic rules and role in neurological pathogenesis. J. Cell Sci. 126:2319–2329.

Gupta, V., K.J. Palmer, P. Spence, A. Hudson, and D.J. Stephens. 2008. Kinesin-1 (uKHC/KIF5B) is required for bidirectional motility of ER exit sites and efficient ER-to-Golgi transport. Traffic. 9:1850–1866.

Ho, W.C., V.J. Allan, G. van Meer, E.G. Berger, and T.E. Kreis. 1989. Reclustering of scattered Golgi elements occurs along microtubules. Eur. J. Cell Biol. 48:250–263.

Hunt, S.D., A.K. Townley, C.M. Danson, P.J. Cullen, and D.J. Stephens. 2013. Microtubule motors mediate endosomal sorting by maintaining functional domain organization. J. Cell Sci. 126:2493–2501.

Lewis, M.J., and H.R. Pelham. 1990. A human homologue of the yeast HDEL receptor. Nature. 348:162–163.

Lippincott-Schwartz, J., N.B. Cole, A. Marotta, P.A. Conrad, and G.S. Bloom. 1995. Kinesin is the motor for microtubule-mediated Golgi-to-ER membrane traffic. J. Cell Biol. 128:293–306.

Lippincott-Schwartz, J., J.G. Donaldson, A. Schweizer, E.G. Berger, H.P. Hauri, L.C. Yuan, and R.D. Klausner. 1990. Microtubule-dependent retrograde transport of proteins into the ER in the presence of brefeldin A suggests an ER recycling pathway. Cell. 60:821–836.

Lippincott-Schwartz, J., L.C. Yuan, J.S. Bonifacino, and R.D. Klausner. 1989. Rapid redistribution of Golgi proteins into the ER in cells treated with brefeldin A: evidence for membrane cycling from Golgi to ER. Cell. 56:801–813.

Mayor, S., J.F. Presley, and F.R. Maxfield. 1993. Sorting of membrane components from endosomes and subsequent recycling to the cell surface occurs by a bulk flow process. J. Cell Biol. 121:1257–1269.

Munro, S., and H.R. Pelham. 1987. A C-terminal signal prevents secretion of luminal ER proteins. Cell. 48:899–907.

Palmer, K.J., H. Hughes, and D.J. Stephens. 2009. Specificity of cytoplasmic dynein subunits in discrete membrane-trafficking steps. Mol. Biol. Cell. 20:2885–2899.

Polishchuk, E.V., A. Di Pentima, A. Luini, and R.S. Polishchuk. 2003. Mechanism of Constitutive Export from the Golgi: Bulk Flow via the Formation, Protrusion, and En Bloc Cleavage of large trans-Golgi Network Tubular Domains. Mol. Biol. Cell. 14:4470–4485.

Presley, J.F., N.B. Cole, T.A. Schroer, K. Hirschberg, K.J. Zaal, and J. Lippincott-Schwartz. 1997. ER-to-Golgi transport visualized in living cells. Nature. 389:81–85.

Roux, A., G. Cappello, J. Cartaud, J. Prost, B. Goud, and P. Bassereau. 2002. A minimal system allowing tubulation with molecular motors pulling on giant liposomes. Proc. Natl. Acad. Sci. USA. 99:5394–5399.

Schindelin, J., I. Arganda-Carreras, E. Frise, V. Kaynig, M. Longair, T. Pietzsch, S. Preibisch, C. Rueden, S. Saalfeld, B. Schmid, J.Y. Tinevez, D.J. White, V. Hartenstein, K. Eliceiri, P. Tomancak, and A. Cardona. 2012. Fiji: an open-source platform for biological-image analysis. Nature Methods. 9:676–682.

Shima, D.T., S.J. Scales, T.E. Kreis, and R. Pepperkok. 1999. Segregation of COPI-rich and anterograde-cargo-rich domains in endoplasmic-reticulum-to-Golgi transport complexes. Curr. Biol. 9:821–824.

Simpson, J.C., T. Nilsson, and R. Pepperkok. 2006. Biogenesis of tubular ER-to-Golgi transport intermediates. Mol. Biol. Cell. 17:723–737.

Stauber, T., J.C. Simpson, R. Pepperkok, and I. Vernos. 2006. A role for kinesin-2 in COPI-dependent recycling between the ER and the Golgi complex. Current biology : CB. 16:2245–2251.

Stephens, D.J. 2012. Functional coupling of microtubules to membranes - implications for membrane structure and dynamics. J. Cell Sci. 125:2795–2804.

Thyberg, J., and S. Moskalewski. 1985. Microtubules and the organization of the Golgi complex. Exp. Cell Res. 159:1–16.

Tomas, M., E. Martinez-Alonso, J. Ballesta, and J.A. Martinez-Menarguez. 2010. Regulation of ER-Golgi intermediate compartment tubulation and mobility by COPI coats, motor proteins and microtubules. Traffic. 11:616–625.

Watson, P., R. Forster, K.J. Palmer, R. Pepperkok, and D.J. Stephens. 2005. Coupling of ER exit to microtubules through direct interaction of COPII with dynactin. Nat. Cell Biol. 7:48–55.

Yadav, S., M.A. Puthenveedu, and A.D. Linstedt. 2012. Golgin160 recruits the Dynein motor to position the Golgi apparatus. Dev. Cell. 23:153–165.

